# Enhanced synaptic transmission in the extended amygdala and altered excitability in an extended amygdala to brainstem circuit in a Dravet syndrome mouse model

**DOI:** 10.1101/2020.09.01.267112

**Authors:** Wen Wei Yan, Maya Xia, Alyssa Levitt, Nicole Hawkins, Jennifer Kearney, Geoffrey T. Swanson, Dane Chetkovich, William P. Nobis

**Author notes:** denotes equal contribution (Wen Wei Yan, Maya Xia). **Corresponding Author:** William P. Nobis, 6130A MRB 3/Bio Sci Building, 465 21st Ave S, Nashville, TN 37235.

## Abstract

**Objective:** Dravet syndrome (DS) is a severe, early-onset epilepsy with an increased incidence of sudden death. Evidence of interictal breathing deficits in DS suggest that alterations in subcortical projections to brainstem nuclei may exist, which might be driving comorbidities in DS. The aim of this study was to determine if a subcortical structure, the bed nucleus of the stria terminalis (BNST) in the extended amygdala, is activated by seizures, exhibits changes in excitability, and expresses any alterations in neurons projecting to a brainstem nucleus associated with respiration, stress response and homeostasis.

**Methods:** Experiments were conducted using F1 mice generated by breeding 129.Scn1a^+/-^ mice with wildtype C57BL/6J mice. Immunohistochemistry was performed to quantify neuronal c-fos activation in DS mice after observed spontaneous seizures. Whole cell patch clamp and current clamp electrophysiology recordings were conducted to evaluate changes in intrinsic and synaptic excitability in the BNST.

**Results:** Spontaneous seizures in DS mice significantly enhanced neuronal c-fos expression in the BNST. Further, the BNST had altered AMPA/NMDA postsynaptic receptor composition and showed changes in spontaneous neurotransmission, with greater excitation and decreased inhibition. BNST to parabrachial nucleus (PBN) projection neurons exhibited intrinsic excitability in wildtype mice, while these projection neurons were hypoexcitable in DS mice.

**Significance:** The findings suggest that there is altered excitability in neurons of the BNST, including BNST to PBN projection neurons, in DS mice. These alterations could potentially be driving comorbid aspects of DS outside of seizures, including respiratory dysfunction and sudden death.

**SIGNIFICANCE STATEMENT:** Dravet syndrome (DS) is an early-onset epilepsy with an increased risk of sudden death. We determined that there are alterations in a subcortical nucleus, the bed nucleus of the stria terminalis (BNST) of the extended amygdala, in a murine DS model. The BNST is involved in stress, anxiety, feeding, and respiratory function. We found enhanced activation in the BNST after seizures and alterations in basal synaptic neurotransmission–with enhanced spontaneous excitatory and decreased spontaneous inhibitory postsynaptic events. Evaluating those neurons that project to the parabrachial nucleus (PBN), a nucleus with multiple homeostatic roles, we found them to be hypoexcitable in DS. Alterations in BNST to brainstem projections could be implicated in comorbid aspects of DS, including respiratory dysfunction and sudden death.

## INTRODUCTION

Dravet syndrome (DS) is a developmental and epileptic encephalopathy with multiple neuropsychiatric comorbidities and an increased risk of sudden death (Gataullina and Dulac, 2017). Most cases arise due to a *de novo* variant in the *SCN1A* gene, encoding the sodium channel α subunit Nav1.1. Seizures, while usually present early and often refractory, are not the only manifestation of the disease. Other common comorbidities can be just as impactful on patients and caregivers and include psychomotor delay, cognitive disability, mood disorders, dysautonomia, nutritional issues, and a significant risk of sudden unexpected death in epilepsy (SUDEP) (Villas et al., 2017).

Murine models of DS recapitulate many aspects of human disease including severe epilepsy, cognitive and social deficits, and sudden death (Kalume et al., 2013; Lopez-Santiago and Isom, 2019). Work using these models has begun to unravel the neurophysiological alterations in DS that may be underpinnings of these comorbidities. Cardiac abnormalities and peri-ictal breathing issues have been noted in patients and murine models, with more recent work implicating respiratory decline in SUDEP in DS mice. While a major hypothesis proposes that spreading depolarization and subsequent brainstem depression may trigger respiratory decline (Bernard, 2015), evidence of interictal breathing deficits in DS patients and mice suggests that direct alterations in brainstem respiratory centers or their subcortical projecting neurons may exist (Kim et al., 2018). Indeed, recent work shows altered excitability in a region critical for respiratory control and chemosensation (Kuo et al., 2019). It is also possible, yet largely unexplored, that alterations in subcortical projections to these brainstem nuclei may cause deficits as well, with potential far reaching consequences that could contribute to comorbidities such as temperature dysregulation, mood disorders, feeding, cardiac and respiratory dysfunction and risk for sudden death. Such alterations may be intrinsic to the subcortical structure or driven by seizures and upstream hyperexcitability.

Despite impressive study of DS mouse models and epilepsy, the neuronal basis for seizures in DS is still debated(Isom, 2014); however, it appears to critically involve forebrain disinhibition (Cheah et al., 2012; Favero et al., 2018; Yuan et al., 2019). Multiple electrophysiological studies demonstrate impaired firing of inhibitory neurons and selective deletion of Scn1a in inhibitory neurons alone is sufficient to cause seizures and premature death (Cheah et al., 2012). Clearly, intrinsic deficits in cortical inhibition are driving hyperexcitability; however, little work has been performed on the electrophysiological changes in downstream subcortical targets (Derera et al., 2017; Kuo et al., 2019).

A recent study found that targeting a subcortical structure in DS mice, a nucleus of the extended amygdala, could have a significant effect on seizure induced apneas and mortality due to SUDEP (Bravo et al., 2020). This important study suggests that alterations in a subcortical circuit may be driving an important aspect of DS respiratory dysfunction during seizures. In the present study, we aimed to determine whether another nucleus of the extended amygdala, the bed nucleus of the stria terminalis (BNST), is activated by seizures in DS animals, whether there exists any synaptic or intrinsic excitability changes in this nucleus, and whether neurons projecting to a brainstem structure important for both respiratory function and also overall homeostasis is altered, which could have implications for mechanisms and treatment of important comorbidities in DS.

## METHODS

### Mouse husbandry and genotyping

Animal care and experimental procedures were approved in accordance with the National Institutes of Health Guide for the Care and Use of Laboratory Animals. Mice were group-housed in a mouse facility under standard laboratory conditions (14-h light/10-h dark) and had access to food and water ad libitum, except during hyperthermia-induced seizure experiments.

Scn1a^tm1Kea^ mice were obtained (Miller et al., 2014). The line has been maintained as a co-isogenic strain by continuous backcrossing of null heterozygotes to 129 (129.Scn1a^+/-^). 129S6/SvEvTac mice (stock #129SVE) were obtained from Taconic (Germantown, NY, USA). For experiments, F1 mice were produced by breeding heterozygous 129.Scn1a^+/-^ mice with wildtype C57BL/6J mice (Jackson Laboratory stock 000664). Mice were ear tagged and tail-biopsied between postnatal day 14 and 20 (P14-20). At ~P21, F1 mice were weaned into holding cages containing four to five mice of the same sex and age for experiments.

### Acute slice preparation

Acute brain slices were prepared from P19-30 heterozygous and wildtype littermates of either gender, in accordance with Institutional Animal Care and Use Committee approved protocols. Mice were decapitated under isoflurane, and their brains were removed quickly and placed in an ice-cold sucrose-rich slicing ACSF containing 85 mM NaCl, 2.5 mM KCl, 1.25 mM NaH_2_PO_4_, 25 mM NaHCO_3_, 75 mM sucrose, 25 mM glucose, 10 M DL-APV, 100 μM kynurenate, 0.5 mM Na L-ascorbate, 0.5 mM CaCl_2_, and 4 mM MgCl_2_. Sucrose-ACSF was oxygenated and equilibrated with 95% O_2_/5% CO_2_. Hemisected coronal slices (300 μm) were prepared using a Leica VT1200S vibratome (Leica Biosystems, Wetzlar Germany). Slices were transferred to a holding chamber containing sucrose-ACSF warmed to 30°C and slowly returned to room temperature over the course of 15-30 min. Slices were then transferred to oxygenated ACSF at room temperature containing 125 mM NaCl, 2.4mM KCl, 1.2 mM NaH_2_PO_4_, 25 mM NaHCO_3_, 25 mM glucose, 2 mM CaCl_2_, and 1 mM MgCl_2_ and maintained under these incubation conditions until recording.

### Electrophysiological recordings

Slices were transferred to a submerged recording chamber continuously perfused at 2.0 mL/min with oxygenated ACSF maintained at 30±2°C. Neurons in the dBNST were identified using infrared differential interference contrast on a Scientifica Slicescope II (East Sussex, UK). Whole-cell patch clamp recordings were performed using borosilicate glass micropipettes with tip resistance between 3-6 MΩ. Signals were acquired using an Axon Multiclamp 700B (Molecular Devices, Sunnyvale, CA). Data were sampled at 10 kHz, low-pass filtered at 3 kHz. Access resistances ranged between 5 and 24 MΩ and were continuously monitored. Changes greater than 20% from the initial value were excluded from data analyses. Series resistance was uncompensated. Data was recorded and analyzed using pClamp 11 (Molecular Devices, Sunnyvale, CA).

Current-clamp was performed using a potassium gluconate-based intracellular (in mM: 135 K-gluconate, 5 NaCl, 2 MgCl_2_, 10 HEPES, 0.6 EGTA, 4 Na_2_ATP, 0.4 Na_2_GTP, pH 7.3, 285–290 mOsm). Input resistance was measured immediately after breaking into the cell and was determined from the peak voltage response to a −5 pA current injection. Following stabilization and measurement of the resting membrane potential, current was injected to hold all cells at a membrane potential of −65 mV, maintaining a common membrane potential to account for intercell variability. Changes in excitability were evaluated by measuring action potential dynamics and the number of action potentials fired at increasing 10 pA current steps (−150 to 150 pA). The action potential threshold and the amount of current required to fire an action potential (rheobase) were assessed through a ramp protocol of 120 pA/1s. Parameters related to AP shape, which included AP height, AP duration at half-maximal height (AP half-width), and time to fire an AP (AP latency), were calculated from the first action potential fired during the V-I plot. Hyperpolarization sag was calculated as the difference between the initial maximal negative membrane potential and the steady-state current after negative current injection. Neuron types were determined based on the classifications of Hammack (Hammack et al., 2007; Rodríguez-Sierra et al., 2013).

For assessment of spontaneous activity, a cesium methanesulfonate-based intracellular (in mM: 135 Cs-methanesulfonate, 10 KCl, 1 MgCl_2_, 0.2 EGTA, 4 MgATP, 0.3 Na_2_GTP, 20 phosphocreatine, 10 QX-314, pH 7.3, 285-290 mOsm) was used. Cells were voltage-clamped at −65 mV to monitor spontaneous excitatory postsynaptic currents (sEPSCs) and held at +10 mV to isolate spontaneous inhibitory postsynaptic currents (sIPSCs). sPSCs were measured in 120-second blocks, excluding <5 pA events. Analysis was performed using MiniAnalysis (Synaptosoft).

For evoked experiments, postsynaptic currents were evoked by electrical stimulation using a monopolar glass electrode placed dorsal to the recorded neuron controlled by a SIU91 constant current isolator (Cygnus Tech, Delaware Water Gap, PA). An internal recording solution (in mM: 95 CsF, 25 CsCl, 10 Cs-HEPES, 10 Cs-EGTA, 2 NaCl, 2 Mg-ATP, 10 QX-314, 5 TEA-Cl, and 5 4-aminopyridine, pH 7.3, 285-292 mOSm) was used. To evaluate evoked EPSCs, picrotoxin (25 μM) was added to the ACSF to block both synaptic and extrasynaptic GABA-A receptors to assess post-synaptic glutamate transmission. To assess putative changes in presynaptic release probability, paired pulse ratio (amplitude of EPSC2/ EPSC1) was recorded at 50 ms interstimulus interval while the cells were voltage clamped at −65 mV. Similar experiments were conducted to assess short term plasticity at GABAergic terminals in the presence of kynurenic acid (3 mM) to block ionotropic glutamate receptors. Since eIPSCs have slower kinetics, a 150 ms interstimulus interval was used and PPR was calculated as delta amplitude of IPSC2 divided by amplitude of IPSC1. NMDA/AMPA ratios were calculated based on the peak AMPAR-mediated EPSC amplitude measured at −70 mV, and the NMDAR-mediated EPSC component measured 40 ms following electrical stimulation at a 40 mV holding potential, respectively. The reversal potential for NMDA receptors is near 0 mV (Misra et al., 2000; Wyeth et al., 2014) and it has been previously determined that at this time-point the AMPA component of the ESPC has completely decayed (Fernandes et al., 2009; Hedrick et al., 2017).

### Hyperthermia-induced seizures

Hyperthermia-induced seizure experiments were conducted in P20-28 Scn1a^+/-^ mice and wildtype littermates at age P18-25 in order to prime animals to have more frequent spontaneous seizures (Hawkins et al., 2017). A rodent temperature regulator (TCAT-2DF, Physitemp Instruments, Inc, Clifton, NJ) connected to a heat lamp and RET-3 rectal temperature probe was used. The rectal probe was inserted, and mice acclimated to the temperature probe for 5 min before the hyperthermia protocol was started. Mouse core body temperature was elevated 0.5°C every 2 min until the onset of the first clonic convulsion with loss of posture or until 42.5°C was reached. Mice that reached 42.5°C were held at temperature for 3 min.

Spontaneous behavioral seizures were assessed 24-72 h after hyperthermia protocol by an observer blinded to genotype. Animals were brought to an isolated room in the lab and kept in their home cage with ad libitum access to food and water during the daylight cycle. After witnessing a tonic-clonic convulsive seizure, the animal that had the seizure and a wildtype littermate were killed one hour after the event. To capture baseline c-fos levels, mice were handled and acclimated each day in an isolated room in the lab for three days, after which the animals were perfused as previously discussed.

### C-fos fluorescent immunohistochemistry

After a DS mouse survived an observed seizure, the DS animal that had the seizure, a DS littermate, and a wildtype littermate, both of which did not have any observed seizures, were euthanized. Because c-fos expression peaks 1-2 hours after stimulus onset (Hsieh and Watanabe, 2000), animals were euthanized 1-2 hours after the experimenter witnessed a seizure.

Mice were deeply anesthetized with isoflurane and transcardially perfused with ice-cold PBS followed by 4% paraformaldehyde (PFA). Brains were removed and post-fixed for 48 h at 4°C. Coronal sections (30 μm) were made using a vibratome (Leica Biosytems, Wetzlar, Germany). Antigen retrieval was performed with 10 mM sodium citrate, pH 9.0 for 10 min at 80°C. Slices were cooled, washed with PBS, and immersed in blocking solution (5% normal goat serum and 0.03% Triton X-100 in PBS) for 1 h at room temperature with gentle agitation. The primary antibody (c-fos abcam, Cambridge, UK) was diluted in blocking solution and applied overnight at 4°C. Sections were then washed three times for 5 mins each and incubated with fluorescently-labeled secondary antibodies (Invitrogen, Carlsbad, CA) in blocking solution for 1 h at room temperature. Sections were washed an additional three times with DAPI included in the final wash. Tissues were mounted on glass slides using PermaFlour (ThermoFisher). Images were acquired on a Leica DFC290 (Leica Biosytems, Wetzlar, Germany). The same acquisition parameters and alterations to brightness and contrast in ImageJ were used across all images within an experiment. Images were thresholded using the striatum image for each genotype. Following thresholding, positive cells within the BNST were manually counted using ImageJ by a blinded reviewer. All numbers are reported as a single averaged value for each dBNST and then averaged for each animal.

### Stereotaxic surgeries

Mice were anesthetized with isoflurane (initial dose 3%; maintenance dose 1.5%) and injected intracranially with red retrobeads (Lumafluor, Raleigh, NC). A targeted microinjection of the retrobeads (100 nL) was made into the lateral parabrachial nucleus (PBN) (coordinates from Bregma, medial/lateral ±1.4 mm, anterior/posterior: −4.9 mm, dorsal/ventral: 3.8 mm) (Paxinos and Franklin, 2019). All injections were unilateral. Mice were treated with 5 mg/kg injections of ketoprofen for 48 h following surgery. Electrophysiological recordings were made from these injected animals 48-72 h after injection.

### Experimental design and statistical analysis

The number of animals used and the number of cells evaluated are noted in the results section for each experiment. Animals of either sex were used during this study and potential sexspecific differences were evaluated. We did not assume a Gaussian distribution; as such all comparisons are nonparametric unpaired t-tests (Mann-Witney U test) used to determine statistical significance. Mean±SEM is provided throughout the text. Scatter plots are shown for all data with mean and SEM graphically depicted. Outliers were removed if values were outside the 95% confidence interval of the mean, it is noted in the text when outliers were removed.

## RESULTS

### Spontaneous seizures induce c-fos expression in the dBNST in DS mice

We first wanted to determine if the BNST was activated by seizures in DS mice. To do this, we analyzed expression patterns of the immediate early gene c-fos following seizures in the dorsal BNST (dBNST). Previous studies consistently show robust c-fos expression in hippocampal regions following seizures; however, nuclei of the extended amygdala have not yet been evaluated (Dutton et al., 2017; Huffman et al., 2018). We monitored naive Scn1a^+/-^ mice for spontaneous seizures in their home cage with wildtype littermates following hyperthermia priming (Figure 1a). None of the wildtype littermates experienced a seizure. Following a spontaneous seizure, there was a significant increase in c-fos positive cells in the dBNST compared to wildtype littermates (Figure 1b,c). In contrast, there was no significant increase in c-fos cells in the dorsal striatum in these animals, a region like the BNST that has stress responsiveness (Perrotti et al., 2004). This increased activation appears to result from seizures, as there was no difference in dBNST c-fos activation at baseline, with just stress of handling (Figure 1b,c).

**Figure 1:**
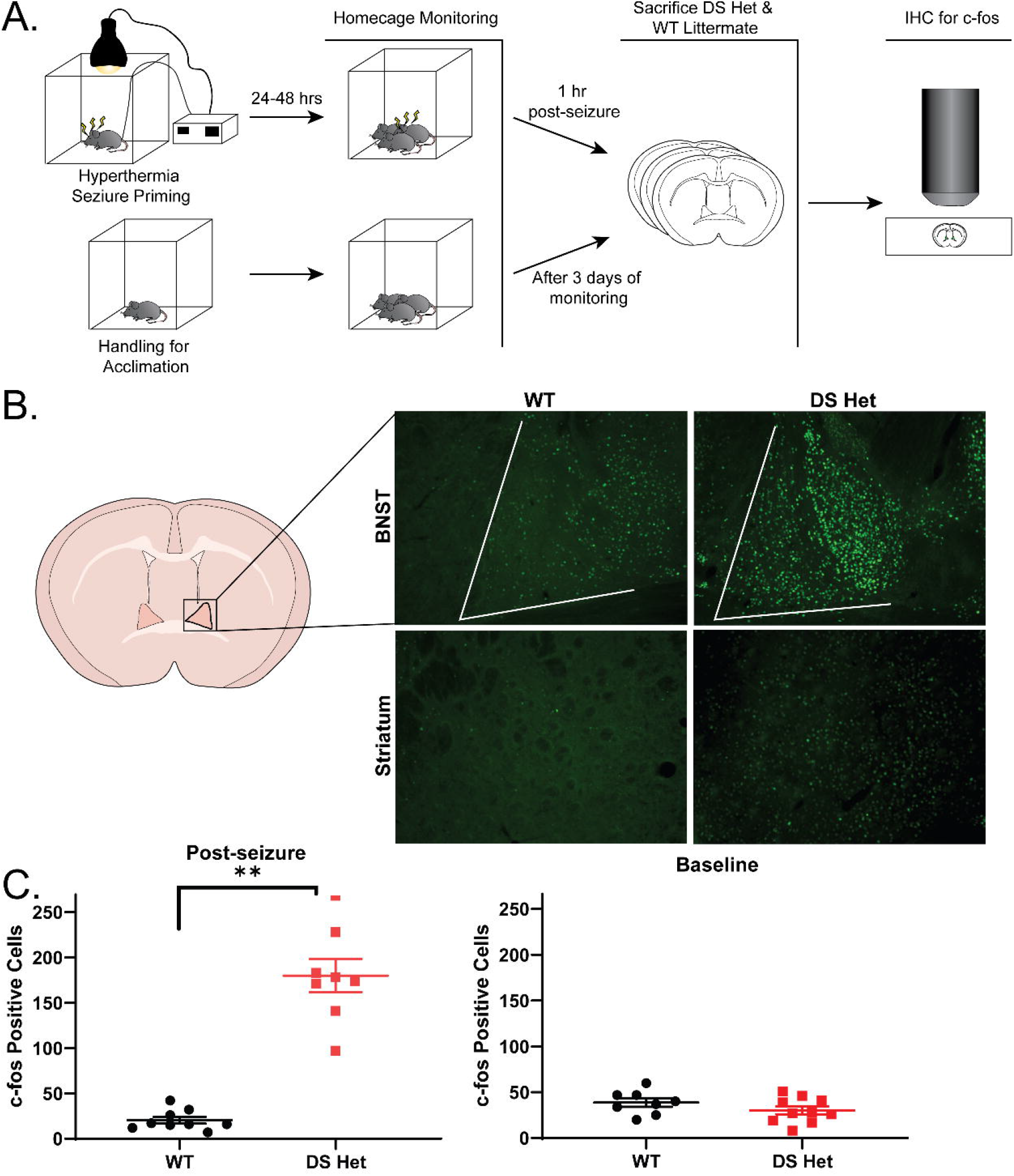
C-fos expression in dBNST increases after spontaneous seizures in DS mice. **A.** For c-fos expression post-seizure, schematic showing timeline of hyperthermia-induced seizure priming, home cage monitoring for seizures, and immunohistochemistry for c-fos in dBNST of DS and WT cage mates. For baseline c-fos levels, timeline of acclimation, home cage monitoring, and immunohistochemistry for c-fos in dBNST. **B**. Illustration of coronal section denotes dBNST region. Representative images of c-fos in dBNST and striatum in DS and WT mice 1 h post-seizure. Striatum served as a negative control for c-fos expression. **C**. Scatter plot showing significant difference in mean number of c-fos activated cells in dBNST sections for WT and DS mice after a spontaneous seizure (WT: n=9, DS: n=8, p < 0.001). No significant difference in c-fos expression in dBNST at baseline levels (WT: n=8, DS: n=10, p=0.26).

### Altered postsynaptic receptor composition in the dBNST in DS mice without changes in release probability

As an initial electrophysiological investigation into potential alterations in dBNST neurons in DS mice, we measured evoked synaptic properties (Figure 2a,b). First, we evaluated synaptic facilitation, a form of short-term plasticity. We bath applied either GABA-A or ionotropic glutamate receptor blockers to isolate excitatory or inhibitory evoked events. We evoked pairs of postsynaptic currents by delivering electrical stimulation, allowing us to measure a paired pulse ratio (PPR). PPR alterations reflect changes in the probability of presynaptic neurotransmitter release (Regehr, 2012). Such alterations are seen in the dentate gyrus in other DS mouse models (Tsai et al., 2015). We found no changes in excitatory (WT: 1.38±0.09 n=9 cells from 6 mice, DS: 1.39±0.9 n=11 cells from 7 mice, p>0.99, Figure 2a) or inhibitory (WT: 0.92±0.08 n=6 cells from 5 mice, DS: 0.99±0.09 n=6 cells from 5 mice, p=0.59, Figure 2b) PPR in the dBNST. We further investigated the contribution of postsynaptic receptors to these evoked excitatory currents. We measured AMPA-receptor mediated EPSCs by voltage clamping the cell at −70 mV and NMDA-receptor mediated currents by holding the same cell at +40 mV. This allowed us to calculate AMPA/NMDA ratio for each cell. Changes in AMPA/NMDA ratio are a potential hallmark for evidence of plasticity and are seen after learning and memory tasks, but also in pathologic situations such as temporal lobe epilepsy models (Amakhin et al., 2017). We found a significant increase in this ratio in DS mice in dBNST neurons (WT: 1.9±0.3 n=7 cells from 5 mice, DS: 3.3±0.4 n=14 cells from 8 mice, p=0.04, Figure 2c).

**Figure 2:**
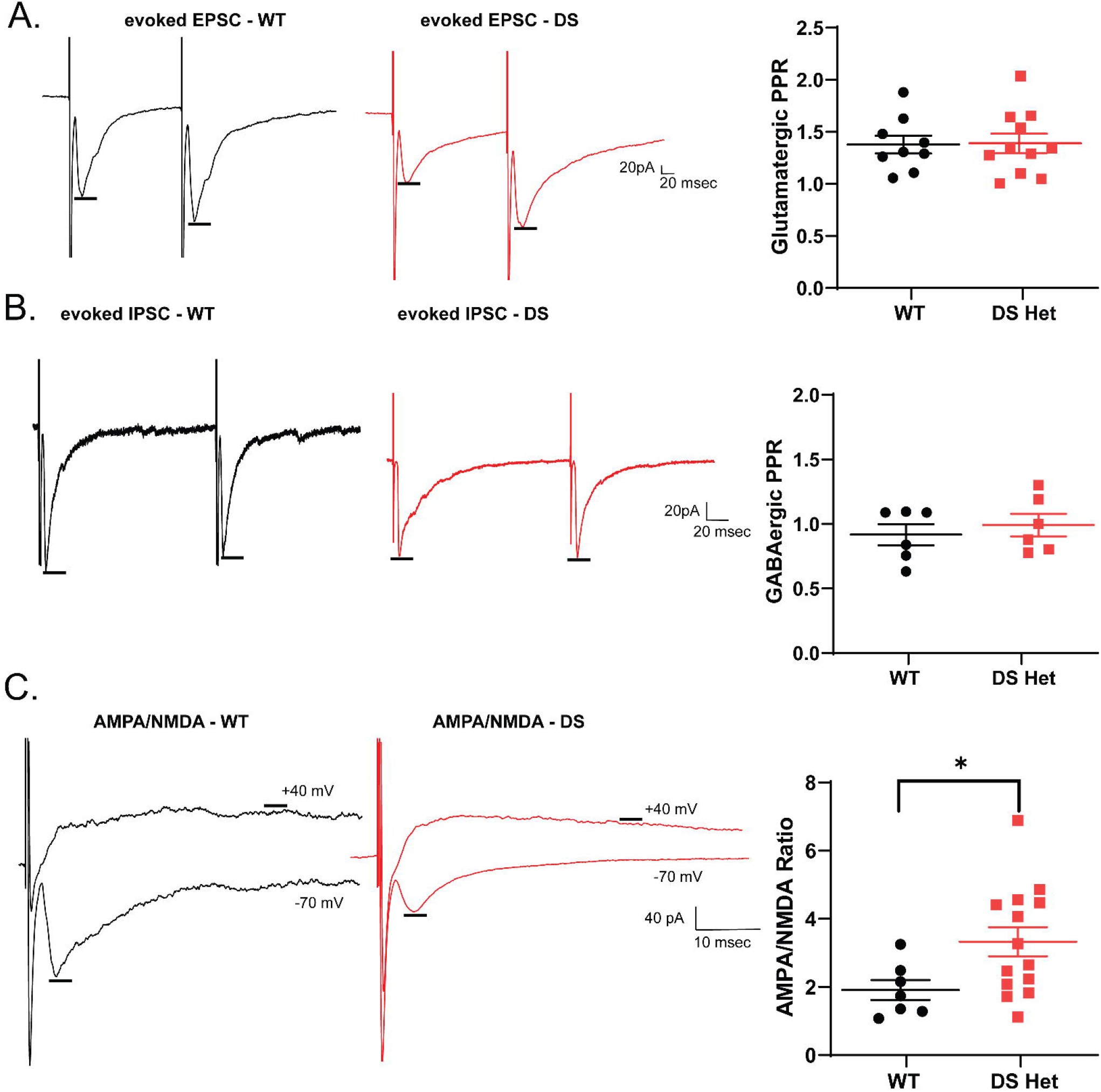
Postsynaptic receptor composition in dBNST is altered in DS mice, but there are no changes in release probability. **A.** Representative evoked EPSC trace of WT (black) and DS (red) neurons in a bath with GABA-A receptor blockers. Scatter plot showing no significant change in excitatory paired pulse ratios (WT: n=9, DS: n=11, p>0.99). **B.** Representative evoked IPSC trace of WT and DS neurons in a bath with ionotropic glutamate receptor blockers. Scatter plot showing no change in inhibitory paired pulse ratios (WT: n=6, DS: n=6, p=0.59). **C.** Representative traces of AMPA-receptor and NMDA-receptor mediated EPSCs in WT and DS neurons. Scatter plot showing significant increase in AMPA/NMDA ratio in DS mice (WT: n=7, DS: n=14, p=0.04).

### Decreased spontaneous inhibitory and enhanced excitatory neurotransmission in DS mice

Excitatory and inhibitory balance is altered in DS mice in cortical and hippocampal regions; to examine the effect of this on the dBNST, we measured both spontaneous excitatory and inhibitory postsynaptic currents. Using a Cs-methanesulfonate internal solution (see methods), we could measure both spontaneous EPSCs when voltage clamped at −65 mV and spontaneous sIPSCs in the same cells when voltage clamped at +10 mV (Figure 3a). While there were no changes in frequency of excitatory events (WT: 3.7±1.9 Hz n=15 cells from 7 mice, DS: 3.9±0.5 Hz n=15 cells from 8 mice, p=0.68, Figure 3c, all experiments in this section are from the same number of mice/cells), there was a significant increase in amplitude of spontaneous events (WT: 13.8±0.77 pA, DS: 17.7±1.25 pA, p=0.016, Figure 3f,g). This is consistent with the increase in AMPA/NMDA, which could result from either a decrease in NMDA synaptic currents or a selective increase in AMPA synaptic currents. Spontaneous inhibitory events in the dBNST were diminished, with a trend for a decrease in sIPSC frequency (WT: 4.8±0.9 Hz n=14 cells from 7 mice with one high outlier removed, DS: 2.8±0.4 Hz, p=0.056, Figure 3c,e).

**Figure 3:**
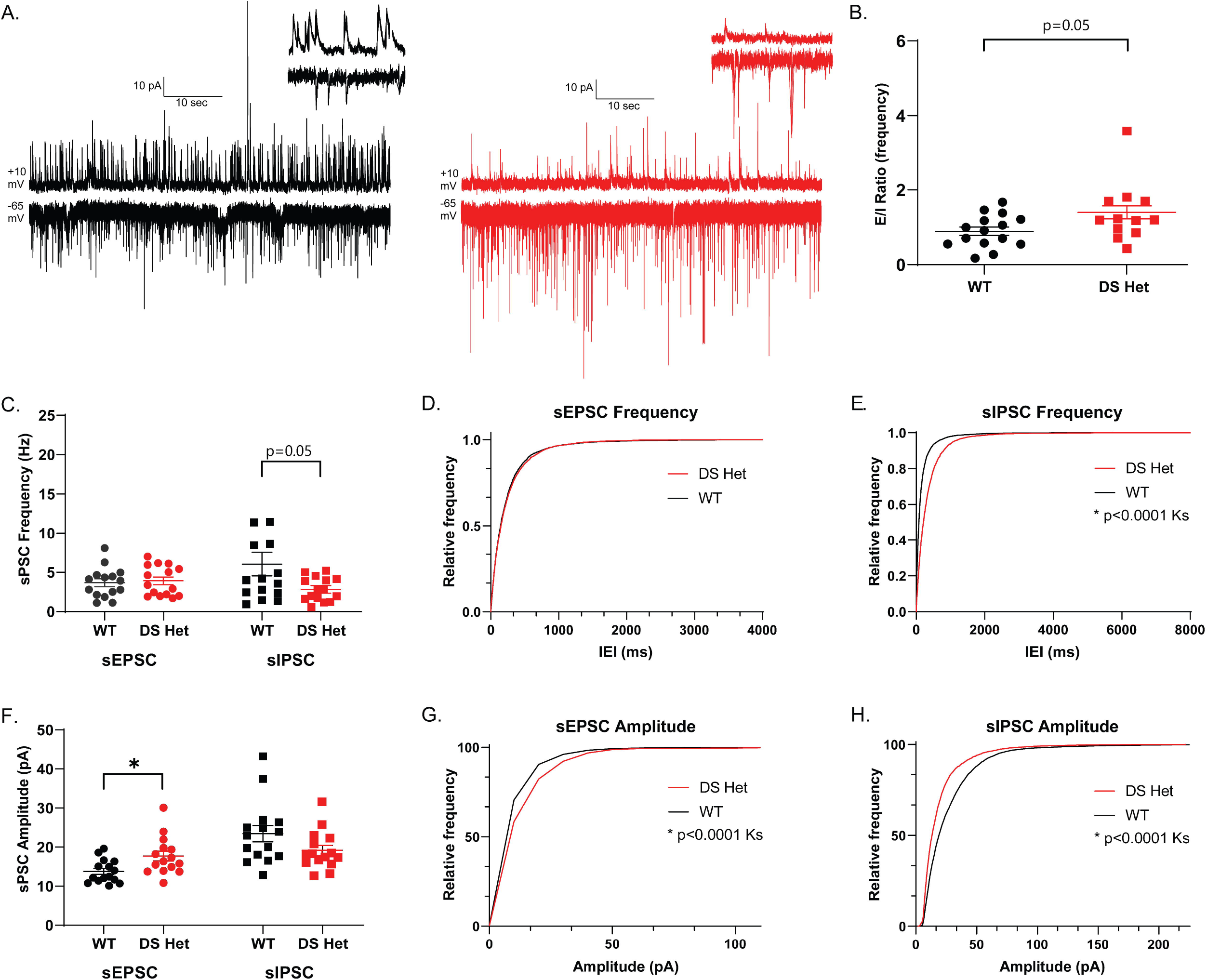
Spontaneous inhibitory neurotransmission is decreased in DS mice, while spontaneous excitatory neurotransmission is enhanced. **A.** Example 60 second traces of WT (black) and DS (red) recordings of spontaneous inhibitory postsynaptic currents (sIPSCs) at +10 mV (top) and spontaneous excitatory postsynaptic currents (sEPSCs) at −65 mV (bottom). Inset of 2 second trace shown (top right). **B.** Scatter plots showing trend for an increase in E/I ratio of frequency of sEPSCs to sIPSCs in DS mice (p=0.059). **C.** Scatter plots showing no significant change in sEPSC frequency (WT: n=15 cells, DS: n=15 cells for all, p=0.68) and trend for a decrease in sIPSC frequency in DS mice (p=0.056). **D, E.** Cumulative probability plots for sEPSC frequency showing no change, while sIPSC frequency showing significant decrease in DS. **F.** Scatter plots showing significant increase in sEPSC amplitude (p=0.016) and a decrease in sIPSC amplitude in DS mice (p=0.09). **G, H.** Cumulative probability plots for sEPSC amplitude showing significant increase, while sIPSC amplitude showing significant decrease in DS.

Given the heterogeneous nature of BNST inputs and intrinsic circuits, a decrease in the number of release sites could be driving this change (Li et al., 2012). There was also a trend for a decrease in the sIPSC amplitude in DS mice (WT: 23.5±2.1 pA, DS: 19.2±1.3 pA, p=0.09, Figure 3f,h). Using this approach to look at both sEPSC and sIPSC within neurons, we calculated an E-I balance for each, determining that there was a trend for an increase in DS mice in frequency of excitatory to inhibitory PSCs (WT: 0.89±0.11, DS: 1.38±0.80 n=13 from 7 mice with two high outliers removed, p=0.059, Figure 3b). Overall, there is an increase in basal synaptic drive in the dBNST in DS mice.

### Minor changes in overall intrinsic excitability of dBNST neurons in DS mice

The NaV1.1 channel is a component of many inhibitory neurons and its loss of function in DS models results in diminished excitability and altered AP dynamics. To investigate whether alterations exist in intrinsic excitability of the largely GABAergic population of both projection and interneurons in the dBNST, we first used brief voltage steps to measure intrinsic properties including input resistance (WT: 625±39.3 MΩ n=18 cells from 9 mice, DS: 666±75.3 MΩ n=19 cells from 8 mice, p=0.96, all experiments in this section are from the same number of mice/cells) and capacitance (WT: 49±4.0 pF, DS: 56±7.0 pF, p=0.62) prior to assessing AP kinetics and excitability in current clamp. After recording the resting membrane potential (RMP) (WT: −60.5±0.56 mV, DS: −60.7±0.73 mV, p=0.85, Figure 4d), all measurements were recorded in neurons clamped at −65 mV to account for the slight variabilities of RMPs between neurons. Overall, there was no change in the number of generated APs across the range of current injections (Figure 4b), nor were there changes in the rheobase (minimum current required to elicit AP) (WT: 57±4.2 pA, DS: 49±4.5 pA, p=0.12, Figure 4e) between the groups. Similarly, AP latency, amplitude and half-width were unchanged between groups (Figure 4h-j). Notably, the threshold to fire the first AP was decreased in DS mice (WT: −32.9±1.28 mV, DS: −39.6±1.72 mV, p=0.007, Figure 4f), indicating the slight increase in excitability. The BNST is a heterogeneous population; however, the neurons can be divided based on their intrinsic membrane currents (Egli and Winder, 2003; Hammack et al., 2007). When subdividing into these categories (Type I, Type II, Type III, and other), there were no differences in intrinsic excitability between WT and DS groups, aside from Type I neurons displaying a lower threshold (WT: −32.6±1.56 mV n=8 cells, DS: −38.1±1.22 mV n=5 cells, p=0.02).

**Figure 4:**
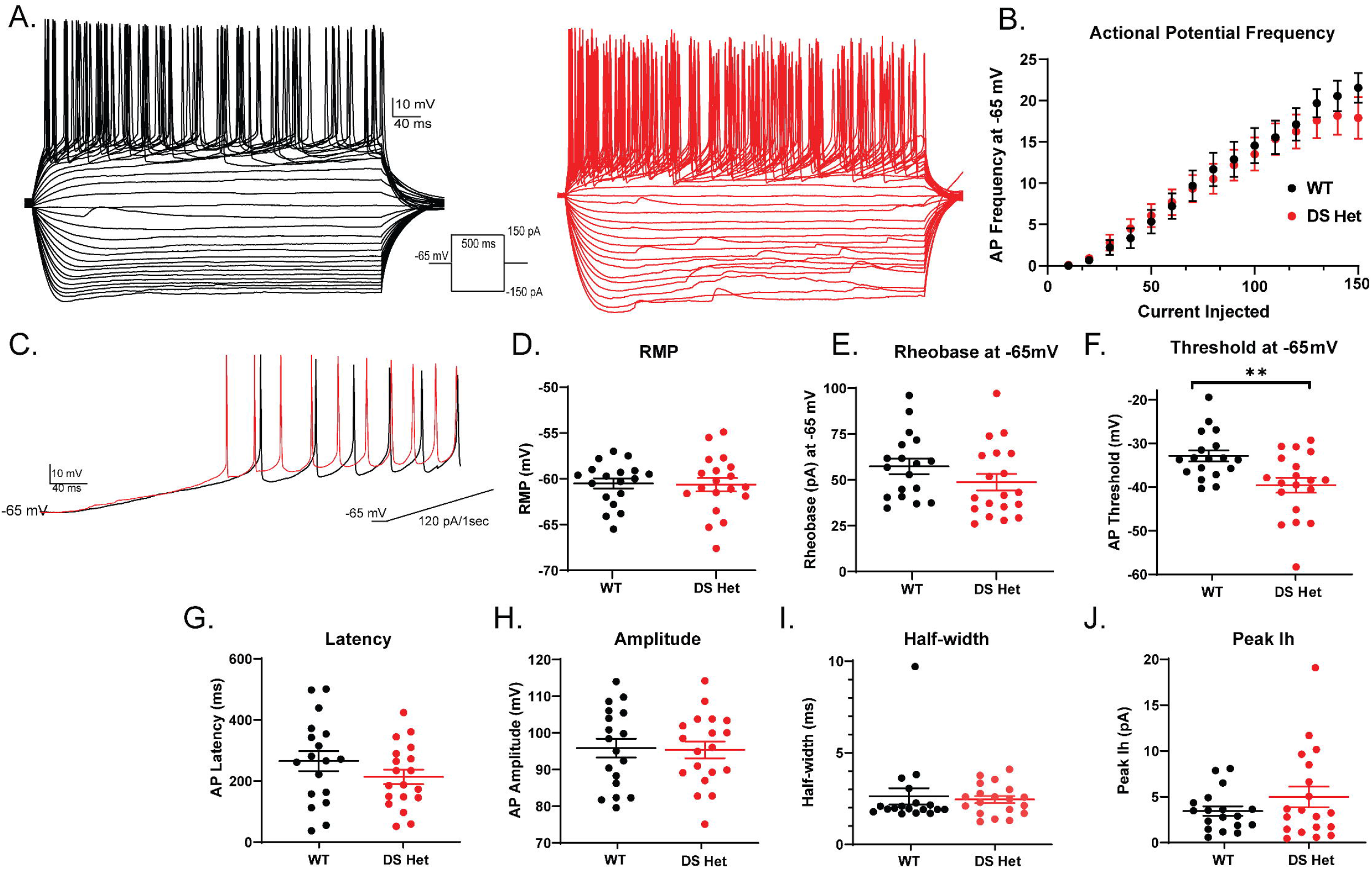
Overall intrinsic excitability of neurons is slightly enhanced in the dBNST of DS mice as a result of lower action potential threshold. **A.** Representative traces of action potentials during current injection protocol in WT (black) and DS (red). **B.** Summary plot showing no difference in action potential frequency after current injection. **C.** Representative trace of ramp current injection in WT (black) and DS (red). **D, E.** Scatter plots showing no difference in resting membrane potential (WT: n = 18, DS: n = 19, p=0.85) and rheobase (p=0.12). **F.** Scatter plot showing significantly lower action potential threshold in DS (p=0.007). **G, H, I, J.** Scatter plots showing no changes in action potential latency, amplitude, half-width, and peak Ih in WT and DS mice.

### dBNST neurons projecting to the PBN are a more excitable population than non-PBN projecting neurons

To determine how changes in the BNST may be influencing downstream targets driving DS comorbidities, we chose to focus on neurons that project to the parabrachial nucleus (PBN). The PBN is a brainstem structure involved in arousal, respiratory function and overall homeostatic functions (Campos et al., 2018). To target these neurons, we stereotaxically injected red retrobeads unilaterally into the PBN and performed whole-cell current clamp experiments from both fluorescent and non-fluorescent neurons (Figure 5a-c) 48-72 h after injection. These neurons on average had a significantly higher input resistance (Fluorescent: 690±60 MΩ n=10 cells from 8 mice, Non-fluorescent: 490±33 MΩ n=8 cells from 8 mice, unpaired t-test p=0.02, Figure 5g, all experiments in this section are from the same number of mice/cells) and a trend for a larger capacitance (Fluorescent: 52±6.7 pF, Non-fluorescent: 34±6.8 pF, unpaired t-test p=0.11), consistent with BNST projection neurons (Luster et al., 2019). Furthermore, we found that dBNST neurons projecting to the PBN are more excitable—firing more APs across ranges of current injections (two-way ANOVA F(15,210)=33.32 p<0.0001 for main effect of group, Figure 5f) and with a significantly lower rheobase (Fluorescent: 48±2.1 pA, Non-fluorescent: 65±5.7 pA, unpaired t-test p=0.009, Figure 5h). Otherwise, there were no differences in RMP, AP kinetics, peak Ih, or threshold in these neurons in wildtype littermates. Previous reports indicate that neurons in the rat and mouse BNST can be categorized into at least three distinct neuronal subtypes based on voltage responses to transient current steps (Egli and Winder, 2003; Hammack et al., 2007). All these physiologically distinct types of neurons were represented in the population of BNST neurons projecting to the PBN, suggesting that changes in excitability in these projection neurons are not simply an enrichment for one type. The plurality of these were Type I neurons (n=5/10 cells) but Type II (n=2/10 cells), Type III (n=1/10 cells) and other (n=2/10) are present. The fact that there is not a strict predominance of a single physiological type in those neurons projecting to the PBN is not unexpected as neither neuronal morphology, expression of neuropeptide markers, nor projection location has previously been shown to associate with a particular physiological sub-type (Ch’ng et al., 2019; Rodríguez-Sierra et al., 2013; Silberman et al., 2013).

**Figure 5:**
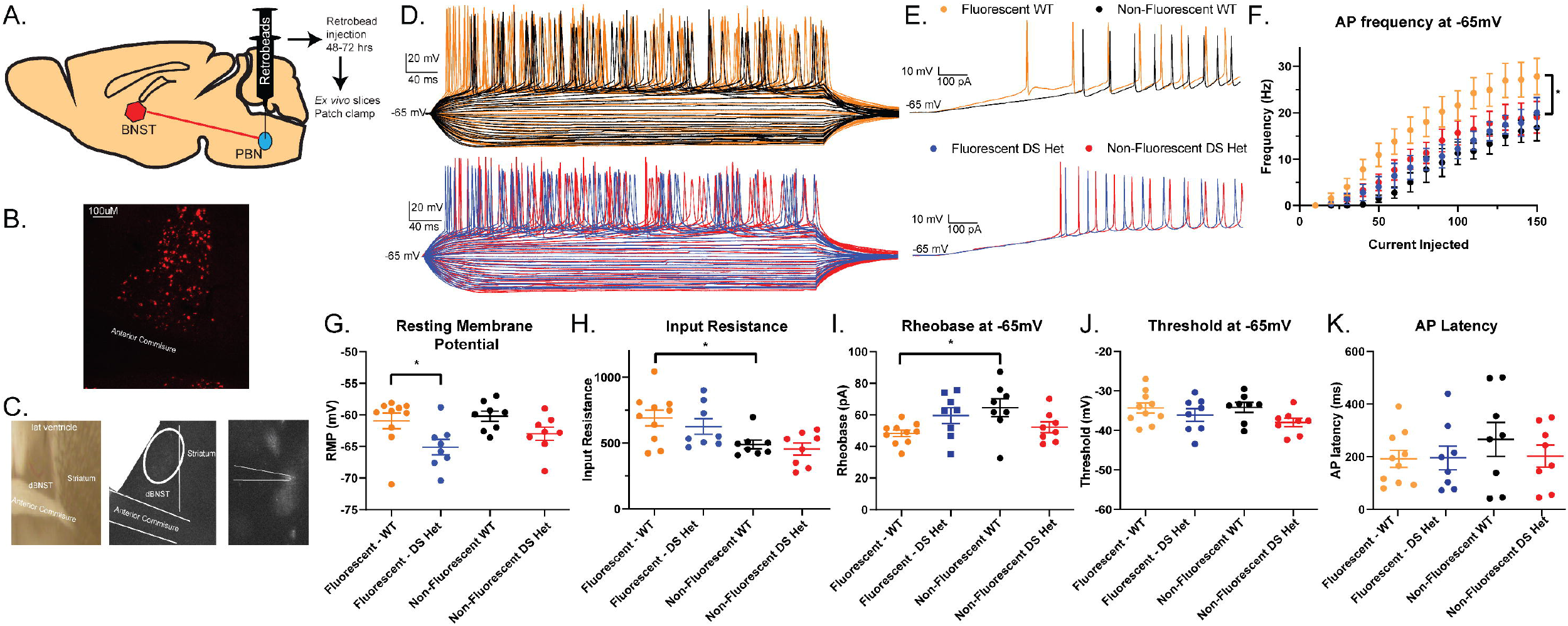
Neurons projecting from the dBNST to PBN are more excitable in WT mice and exhibit hypoexcitability in DS mice. **A.** Schematic showing unilateral retrobead injection into PBN. **B.** Representative image of fluorescent neurons in dBNST following retrobead injection. **C.** Representative image of fluorescent neurons being identified for patch clamp. **D.** Representative traces of action potentials during current injection protocol in WT fluorescent neurons (top, orange), WT non-fluorescent (black), DS fluorescent (bottom, blue), and DS non-fluorescent (red). **E.** Representative trace of ramp current injection in WT (top) and DS (bottom). **F.** Summary plot showing dBNST to PBN projection neurons in WT firing more action potentials and no change in DS neurons across range of current injections (p<0.0001 two-way ANOVA). **G.** Scatter plot showing significant decrease in resting membrane potential of dBNST to PBN projection neurons in DS compared to WT (Fluorescent WT: n=10, Non-fluorescent WT: n=8, Fluorescent DS: n=8, Non-fluorescent DS: n=8, p=0.05 Tukey post-hoc). **H.** Scatter plot showing significant increase in membrane resistance in PBN projecting neurons compared to other neurons in WT (p=0.02) **I.** Scatter plot showing significant decrease in rheobase in fluorescent neurons compared to non-fluorescent neurons in WT (p=0.009). **J, K.** Scatter plots showing no significant changes in threshold and AP latency.

### dBNST neurons projecting to the PBN are hypoexcitable in DS mice

To determine how this potentially important projection neuron was altered in DS mice, we similarly performed recordings from dBNST neurons in DS mice unilaterally injected with red retrobeads in the PBN. Fluorescent neurons in DS mice had a significantly hyperpolarized RMP compared to fluorescent neurons in WT mice (Fluorescent WT: −61 ±1.2 mV n=10 cells from 8 mice, Non-fluorescent WT: −60±0.82 mV n=8 cells from 8 mice, Fluorescent DS: −65±1.2 mV n=8 cells from 6 mice, Non-fluorescent DS: −63±1.0 n=8 cells from 6 mice, p=0.05 Tukey post-hoc, Figure 5g) and strikingly no longer had any change between non-fluorescent neurons from WT or DS mice in AP fired with respect to current injections (two-way ANOVA Tukey post-hoc, Fluorescent DS vs. Non-fluorescent WT p=0.21, Fluorescent DS vs Non-fluorescent DS p=0.74 for main effect of group, Figure 5) or rheobase. Other parameters including AP amplitude, halfwidth, latency, and peak Ih current were unchanged (Figure 5i-k). In DS mice, dBNST projection neurons are therefore hypoexcitable (hyperpolarized, decreased AP firing) with normal AP kinetics.

## DISCUSSION

The current study was motivated by the relative paucity of investigation of subcortical structures in DS despite the impact that pathology in these regions may have on multiple aspects of DS including sudden death. We utilized the Scn1a^+/-^ Dravet model to perform a detailed electrophysiological examination of the dorsal BNST of the extended amygdala, as it receives inputs from cortical and hippocampal regions (Reichard et al., 2016) and has dense outputs to brainstem structures (Dong and Swanson, 2006). This report shows that the dBNST is activated by seizures in DS mice and that there is marked excitatory-inhibitory synaptic imbalance in this region as well as alterations in postsynaptic receptor composition. We further investigated those BNST neurons projecting to the lateral PBN region in the pons and found that these neurons are a physiologically distinct population that are hypoexcitable in DS mice. These results provide evidence for distinct neuroadaptations in the BNST in DS mice and are complementary to the growing literature investigating the importance of subcortical structures in Dravet (Bravo et al., 2020; Kuo et al., 2019).

### Enhanced synaptic drive and synaptic strength in the BNST in DS mice

We provide evidence of enhanced synaptic drive in the dorsal BNST in DS mice, with an increase in amplitude of spontaneous excitatory events and a decrease in both amplitude and frequency of inhibitory events. We additionally show evidence of postsynaptic changes, with an increase in AMPA/NMDA ratio, which suggests that alterations in postsynaptic glutamate receptors may be driving the increase in amplitude of sEPSCs. This may be driven by an enhanced excitatory drive from cortical pyramidal neurons and the ventral hippocampus, which have increased firing in DS mouse models (Mistry et al., 2014; Tai et al., 2014; Yu et al., 2006) and both project to the BNST. A decrease in inhibitory drive, both amplitude and frequency of events, could be a counterbalancing effect for the enhanced excitatory drive. Alternatively, neuronal firing of inhibitory inputs could also be diminished. The BNST receives the majority of its inhibitory input from the central amygdala (CeA) (Cassell et al., 1999), and unlike the BNST, the CeA has a large population of PV neurons (Wang et al., 2016), which likely have impaired firing in DS mice (Tai et al., 2014; Yu et al., 2006). Finally, our data strongly suggest that the BNST is highly activated following seizures in DS mice. The frequency of seizures and subsequent strong neuronal activation of this region might be driving alterations in AMPA/NMDA ratio and producing compensatory changes in inhibitory synaptic drive.

### Subtle increase in intrinsic excitability in BNST neurons

We found subtle changes in intrinsic excitability overall, with an increase in intrinsic excitability of dBNST neurons associated with a lowering of threshold to fire an AP. This was, however, not associated with any increase in spike frequency with current injections or other changes in excitability or AP kinetics. The somewhat contradictory nature of this is likely due to the heterogeneity of the BNST neurons and the inability to distinguish between interneurons or projection neurons when doing blind-patching experiments-there may be a population within the dBNST that has a more striking enhancement in intrinsic excitability if they were to be targeted directly. Much of the dBNST neurons are GABAergic, with both interneurons and inhibitory projection neurons. In other brain regions in DS models, inhibitory neurons, particularly PV interneurons, have greatly diminished excitability and changes to AP kinetics. This was not seen overall in the BNST, likely due to the very small to non-existent population of PV interneurons in this nucleus as previously reported (Nguyen et al., 2016) and confirmed (Supplementary figure 1) which was unlikely to be sufficiently sampled by blind-patching.

### Decreased excitability in BNST to PBN projection neurons

We were interested in how any changes in the BNST excitability could alter autonomic functioning, breathing and homeostatic behaviors that are perturbed in DS, so we evaluated dBNST neurons that project to the lateral PBN by using a retrograde tracing approach. We found that these projection neurons are a more excitable population compared to the dBNST as a whole—with a higher input resistance, a lower rheobase, a lower latency to fire APs, and a higher frequency of firing APs with each current injection. These attributes are no longer present in PBN-projecting neurons in DS mice, where these neurons also show a significant hyperpolarizing change in resting membrane potential, implying potential changes in potassium channel currents. These changes may be compensatory and driven by the aforementioned E-I balance increase, or by the epilepsy state itself. Potentially, GABAergic projection neurons to the PBN may have reduced excitability due to loss of NaV1.1 channels (Kalume et al., 2007), however, this seems less likely given the low basal level of scn1a expression (Supplemental Figure 1). These projection neurons may be preferentially bombarded by repeated cortical seizures, producing changes in intrinsic excitability to counteract this effect and overall diminishing output to the PBN.

### Functional Implications

The BNST is a limbic structure implicated in mediating behavioral responses to anxiety and stress and is intimately interconnected to the larger amygdala circuit, hippocampus, and cortical regions (Knight and Depue, 2019; Lebow and Chen, 2016). The BNST may function as an integrator of these higher order inputs, affecting its function over the autonomic nervous system, breathing, stress response and homeostasis through dense projections to hypothalamic, midbrain and brainstem nuclei including the periventricular nucleus, ventral tegmental area, periaqueductal gray, nucleus tractus solitarius, PBN and serotonergic neurons in the dorsal and midbrain raphe (Cassell et al., 1999; Dong and Swanson, 2006, 2004; Pollak Dorocic et al., 2014). In particular, there is evidence that the extended amygdala region may be critically involved in respiratory dysfunction seen during seizures in DS (Kim et al., 2018; Kuo et al., 2019). Electrical stimulation of the human amygdala reliably produces apneas (Dlouhy et al., 2015; Lacuey et al., 2019; Nobis et al., 2019, 2018). This is likely related to stimulation of structures of the extended amygdala rather than the lateral regions (Nobis et al., 2018; Rhone et al., 2020). Similarly, seizure spread to these regions may be driving ictal apneas (Dlouhy et al., 2015; Nobis et al., 2019). The involvement of the extended amygdalar regions (CeA and BNST) in ictal apneas may explain why the amygdala at large does not always produce apneas when activated by seizures (Park et al., 2020). The PBN has a known role in modulation of breathing (Dutschmann and Herbert, 2006; Navarrete-Opazo et al., 2020) and has been targeted to rescue respiratory dysfunction in developmental disorders (Abdala et al., 2016). The BNST input to the PBN can influence respiration (Kim et al., 2013), so it is possible that decreased intrinsic excitability of BNST projection neurons may produce compensatory changes in the PBN that allow strong stimuli in the form of a seizure to now produce an aberrant response, resulting in apneas. This could work in concert with dysfunction in other brainstem respiratory regions such as the retrotrapezoid nucleus (Kuo et al., 2019), which is more involved in setting respiratory rhythm (Guyenet et al., 2019), with disastrous outcomes leading to ictal breathing suppression and SUDEP. The known role of the lateral PBN in cardiovascular function may also be playing a role (Davern, 2014).

Outside of respiratory and cardiovascular function, the BNST inputs to the PBN have been implicated in anxiety, morphine withdrawal, feeding and aversion (Luster et al., 2019; Mazzone et al., 2018; Wang et al., 2019). The PBN itself has important roles in other homeostatic functions including arousal and temperature regulation (Campos et al., 2018). Altered extended amygdalar inputs to this critical brainstem region could be driving some other persistent aspects of DS outside of seizures. This highlights the general need to explore subcortical structures in epilepsy models, in part to shed light on comorbidities of epilepsy states. The few studies that have been performed evaluating these regions in human epilepsy patients show widespread alterations in subcortical network activity including the brainstem that correlates with impairments in vigilance and arousal (Englot et al., 2020). This current study emphasizes the need to evaluate these structures in DS and other epilepsy models, particularly moving to link circuit dysfunction to other behavioral and physiologic comorbidities seen in epilepsy.

In sum, our results show that the BNST of the extended amygdala is activated in Scn1a^+/-^ mice, and that there are neuroadaptations in this region in DS mice. These adaptations include decreased intrinsic excitability of those neurons projecting to the PBN in the brainstem. These alterations could potentially be driving comorbid aspects of DS outside of seizures, including respiratory dysfunction and sudden death.

## Acknowledgements

Thanks to Ye Han and Isabelle Reinteria for technical assistance and support.

## Funding

This work was supported by grants to W.N. from the American Epilepsy Society (Junior Investigator Award), a pilot grant from the National Institute for Neurologic Disease and Stroke’s Center for SUDEP Research (CSR), and the Vanderbilt Faculty Research Scholars (VFRS) award.

## Conflict of Interest

Authors report no conflict of interest.

## Author contribution statement

WY, MX, AL, and WN Performed Research; WN and MX Wrote the paper; NH, JK, GS, and DC Contributed unpublished reagents/analytic tools; WN Designed research; WN, MX, WY and AL Analyzed data

## Notes

### Competing Interest Statement

The authors have declared no competing interest.

