## Supplementary figures and images for "Enhanced synaptic transmission in the extended amygdala and altered excitability in an extended amygdala to brainstem circuit in a Dravet syndrome mouse model"

### Supplemental Figure 1

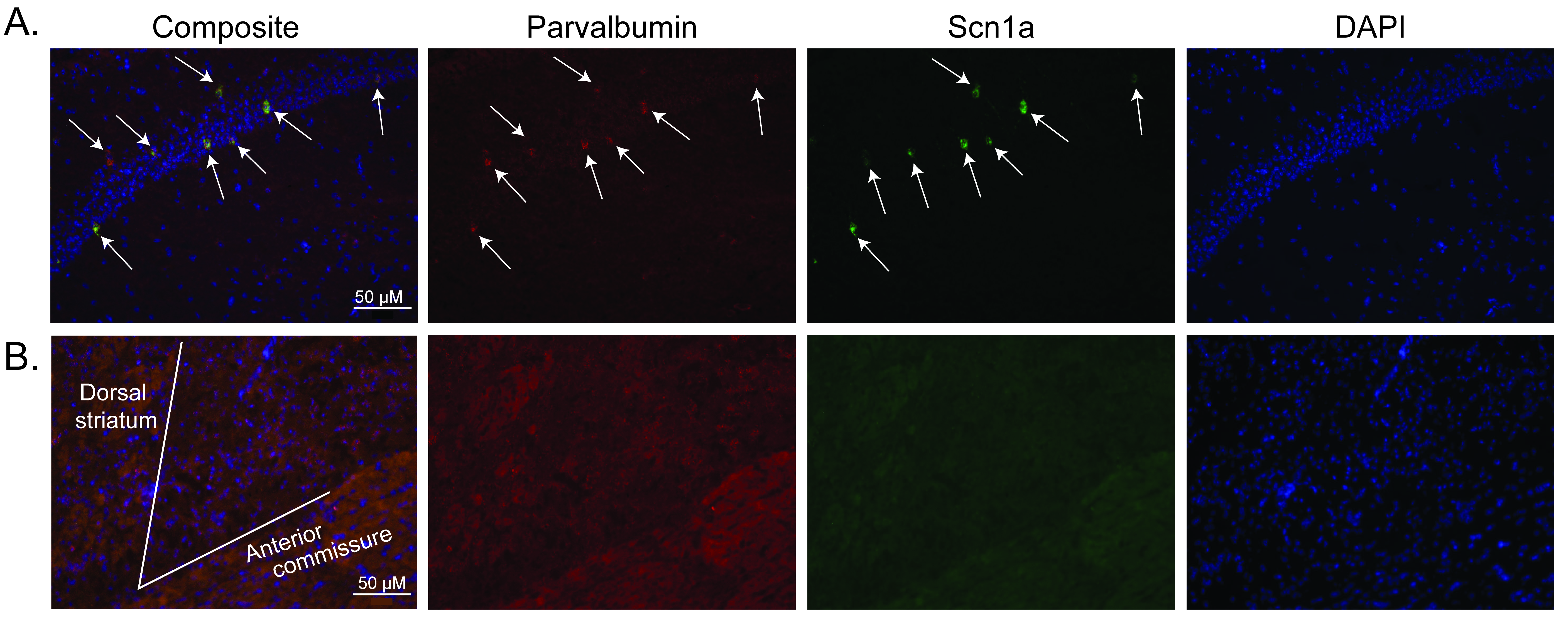
